# Phenome-wide investigation of the causal associations between childhood BMI and adult outcomes: A two-sample Mendelian randomization study

**DOI:** 10.1101/2020.06.01.127530

**Authors:** Shan-Shan Dong, Kun Zhang, Yan Guo, Jing-Miao Ding, Jun-Cheng Feng, Shi Yao, Ruo-Han Hao, Yu Rong, Feng Jiang, Jia-Bin Chen, Hao Wu, Xiao-Feng Chen, Tie-Lin Yang

**Author notes:** Corresponding author: Tie-Lin Yang, Ph.D., Key Laboratory of Biomedical Information Engineering of Ministry of Education, School of Life Science and Technology, Xi’an Jiaotong University, Xi’an 710049, P. R. China.

## Abstract

**Background:** Compelling observational studies have reported that childhood obesity is associated with the risk of many complex diseases in adulthood. However, results from observational studies are unable to fully account for confounding factors. The causal effects of childhood obesity have not been systematically characterized. We aimed to assess the causal associations between childhood body mass index (BMI) and various adult traits/diseases using two-sample Mendelian randomization (MR).

**Methods and findings:** Over 5,000 datasets for adult outcomes were obtained from various resources. After data filtering, 269 adult traits genetically correlated with childhood BMI (*P* < 0.05) were subjected to MR analyses. The number of independent outcomes was 148, setting the significant threshold as *P* < 3.38 × 10^−4^. Inverse-variance weighted method, MR-Egger, weighted median method, and weighted mode method were used to estimate the causal effects.

We identified potential causal effects of childhood obesity on 60 adult traits (27 disease-related traits, 27 lifestyle factors, and 6 other traits). Higher childhood BMI was associated with a reduced overall health rating (β = −0.10, 95% CI: −0.13 to −0.07, *P* = 6.26 × 10^−11^). Findings on diseases included some novel effects, such as the adverse effects of higher childhood BMI on cholelithiasis (OR = 1.26, 95% CI: 1.18 to 1.35, *P* = 3.29 × 10^−5^). For dietary habits, we found that higher childhood BMI was positively associated adult diet portion size (β = 0.26, 95% CI: 0.18 to 0.34, *P* = 7.34 × 10^−11^). Different from the conventional impression, our results showed that higher childhood BMI was positively associated with low calorie density food intake. With 76 adult BMI single-nucleotide polymorphisms (SNPs) as instruments, we confirmed that adulthood BMI was positively associated with heel bone mineral density. However, the association no longer present after excluding the SNPs existing in or in linkage disequilibrium (LD) with childhood BMI. Network MR analyses suggested that past tobacco smoking and portion size mediated 6.39% and 10.90% of the associations between childhood BMI and type 2 diabetes, respectively. The main study limitation is that it is difficult to tease out the independent effects of childhood BMI due to the strong correlation between childhood and adulthood BMI.

**Conclusions:** In summary, we provided a phenome-wide view of the effects of childhood BMI on adult traits. Our results highlight the need to intervene in childhood to reduce obesity from a young age and its later-life effects.

**Author summary:** Why was this study done?

1. Childhood obesity is a worldwide public health problem. The prevalence has increased at an alarming rate.
2. Observational epidemiological studies have reported that childhood obesity is associated with the risk of many complex diseases in adulthood, such as coronary artery disease (CAD) and diabetes. However, observational studies are limited in explaining causality because of possible bias from unmeasured confounding factors.

What did the researchers do and find?

1. A Mendelian randomization (MR) approach was used to provide a phenome-wide view of the causal associations between childhood BMI and adult outcomes. Potential causal effects of childhood obesity on 60 adult traits were identified.
2. Higher childhood BMI was associated with reduced overall health rating, and caused increased risk of some diseases, such as cholelithiasis, hypothyroidism, CAD, and type 2 diabetes (T2D). In contrast, childhood BMI was positively associated with adult heel bone mineral density and low calorie density food intake.
3. Portion size and smoking behavior might mediate the association between childhood BMI and T2D risk.

What do these findings mean?

1. Our results highlight the importance to address rising childhood obesity prevalence rate and early interventions on obesity might help to promote health equity in later life.

## Introduction

Obesity is a worldwide health problem. The prevalence of adult obesity has increased dramatically since the 1980s [1]. It is particularly worrisome that the rate of increase in childhood obesity has been nearly doubled that in adults [1]. Childhood overweight and obesity often persist in adulthood, which increases the risks of premature mortality and physical morbidity across the lifespan [2]. Even more disconcerting is that childhood obesity was associated with an increased risk of multiple comorbidities in adulthood even if the obesity did not persist [3].

Compelling observational studies have reported that childhood obesity is associated with the risk of many complex diseases in adulthood, such as coronary artery disease (CAD) [4], cancers [5], diabetes [6], and polycystic ovary syndrome symptoms [7]. However, results from observational studies are unable to fully account for confounding factors (e.g., socioeconomic status). Therefore, whether the relationship is causal is uncertain.

Mendelian randomization (MR), which uses genetic markers of the exposure as instruments, is now widely used to assess the causal relationship between exposure and outcome [8]. As shown in Fig 1A, MR must satisfy three assumptions [8]: 1) the selected instruments must be associated with the exposure; 2) the instruments must not be associated with confounding factors; 3) the instruments must influence the outcome only through the exposure (no horizontal pleiotropy exists). Conventionally, one-sample MR could be performed by using the two-stage least squares analysis method. Recently, two-sample MR analysis methods using summary-level GWAS data have been developed.[9] With a large amount of GWAS summary data deposited in public databases, two-sample MR analysis provides a cost-efficient way to investigate the potential causal effects of childhood obesity on adult traits. Using this method, previous studies have demonstrated the causal adverse effects of childhood body mass index (BMI) on adult cardiometabolic diseases [10] and osteoarthritis [11]. Using SNPs associated with adult BMI as instruments, two recent MR phenome-wide association studies [12,13] have shown the causal effects of adult obesity on many other traits/diseases. The causal effects of childhood obesity are suspected but have not been systematically characterized.

**Fig 1.**
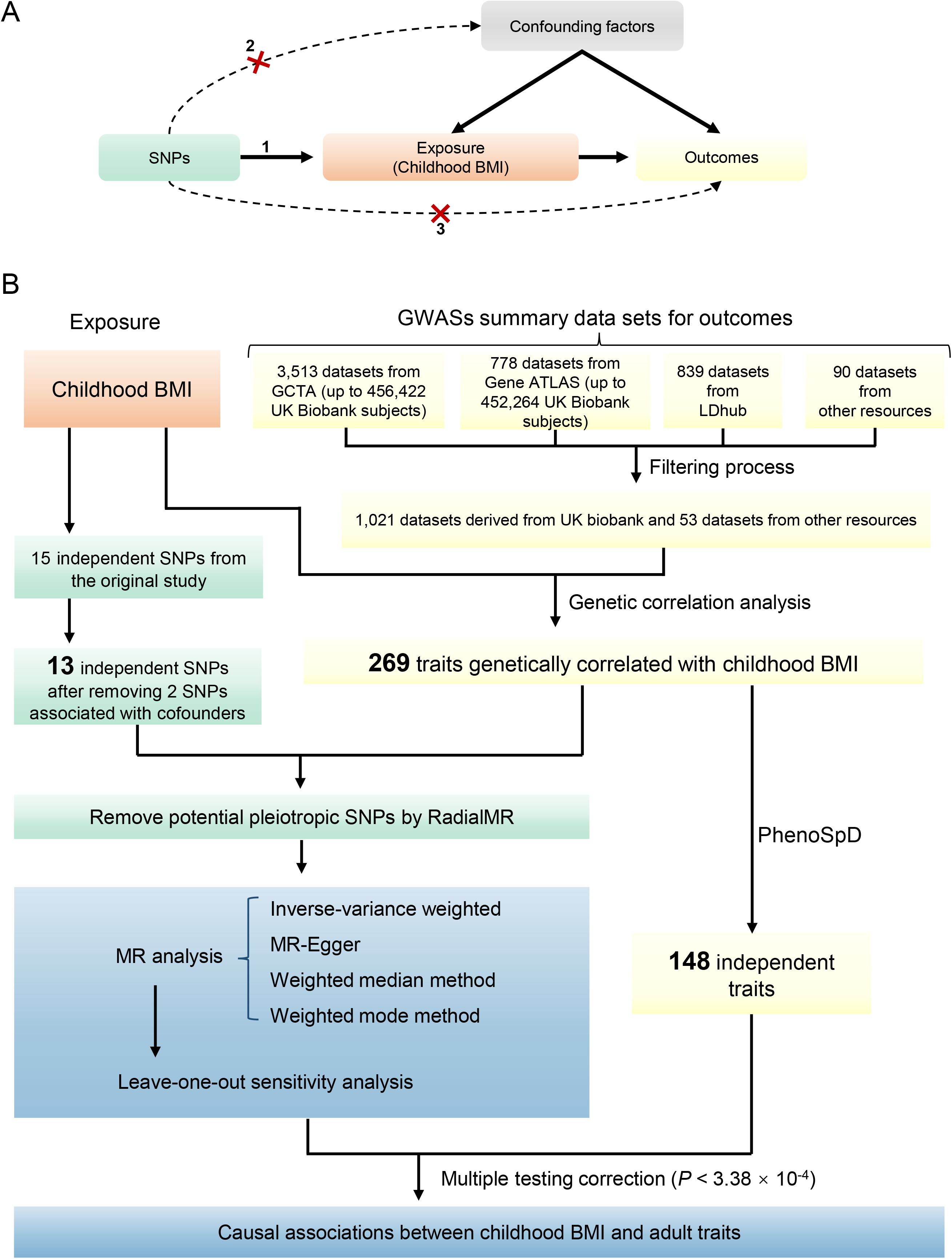
A. Schematic diagram of an MR analysis. Since genetic alleles are independently segregated and randomly assigned, SNPs are not associated with confounding factors that may bias estimates from observational studies. Three assumptions of MR are as follows: 1) the selected instrument is predictive of the exposure; 2) the instrument is independent of confounding factors; 3) there is no horizontal pleiotropy (the instrument is associated with the outcome only through the exposure). B. The analysis pipeline of the current study.

In this study, we performed a MR phenome-wide association study to assess the causal effects of childhood BMI on adult traits/diseases using 2-sample MR with current available GWAS summary data (data collected before May 2019). Our results may offer a systemic view of the causal effects of childhood BMI on adult traits, highlight the importance of intervening early, and provide early clues as to which lifestyle modifications might be helpful for reducing type 2 diabetes (T2D) risk.

## Methods

The outline of the experimental approach used in this study is shown in Fig 1B. The STROBE-MR checklist (https://peerj.com/preprints/27857/) [14] was used to help report the work.

### Summary data resources

#### Childhood BMI

GWAS summary data of childhood BMI were downloaded from the Early Growth Genetics (EGG) consortium (http://egg-consortium.org). The phenotype used in this GWAS was sex- and age-adjusted standard deviation scores of childhood BMI at the latest time point (oldest age) between 2 and 10 years [15]. The GWAS included 47,541 European children in total.

#### Adulthood outcomes

GWAS summary data were mainly obtained from the following resources: 1) 3,513 GWAS summary data on up to 456,422 array-genotyped and imputed UK Biobank individuals (aged between 40 and 69 at recruitment) from the Genome-wide Complex Trait Analysis (GCTA) website; 2) 778 GWAS summary datasets for up to 452,264 UK Biobank individuals from the Gene ATLAS database (http://geneatlas.roslin.ed.ac.uk/); 3) 839 GWAS datasets from the LDhub GWAShare Center (http://ldsc.broadinstitute.org/); 4) 90 datasets from various other resources. All datasets were collected before May, 2019.

Next, we filtered the GWAS summary datasets first using the following criteria:

1. N > 50,000 and both cases and controls are > 10,000 for binary phenotypes.
2. GWAS is based on European population or > 80% of the samples are European.
3. Exclude sex-specific GWAS, unless the trait is only available for a specific sex (e.g., breast cancer).
4. Exclude adolescent traits, parent or sibling traits (e.g., illnesses of father). We also removed traits about adult obesity, since 11 of the 15 childhood BMI SNPs are in LD with adult BMI variants.
5. For one trait with multiple datasets, only the study with the greatest number of subjects was remained.

Finally, in addition to a total of 1,021 datasets derived only from the UK Biobank population, 53 datasets (S1 Table) from other resources remained. All outcomes were recoded to make sure the variables followed increasing patterns. For example, overall health rating was originally coded from 1 to 4 to refer excellent, good, fair, and poor, respectively. Under such situation, an allele positively associated with this trait is actually a risk factor of overall health rating. To avoid misunderstanding, we recoded these traits by changing the plus or minus sign of the beta value in the association results.

### Estimated standardized effect size of SNPs

To enable comparison of effect sizes across studies, we obtained the estimated standardized effect size (β) and standard error (se) as a function of minor allele frequency and sample size as described previously [16] using the following equation:

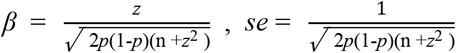

where *z* can be calculated as β/se from the original summary data, *p* is the minor allele frequency, and *n* is the total sample size.

### Genetic correlation analyses

If two traits are causally related and both of them have non-zero heritability, there should be non-zero genetic correlations between them [17]. Therefore, before MR analyses, we firstly carried out genetic correlation analyses using the cross-trait LD Score regression method [18]. All traits that showed nominal genetic correlation with childhood BMI (*P* < 0.05) were classified into three main categories—lifestyle factors, disease-related traits, and others. All disease related traits were further classified according to the International Classification of Diseases 11th Revision (ICD-11). MR analyses were subsequently performed to infer the causal effects of childhood BMI on these outcomes.

### Instruments selection

Fifteen independent SNPs with *P* < 5 × 10^−8^ identified from the original GWAS study [15] for childhood BMI were used as instruments (S2 Table). The genetic risk score of these SNPs explained 2.0% of the variance in childhood BMI [15]. To avoid potential cofounding, we looked up each instrument SNP and their proxies (r^2^ > 0.8) in the PhenoScanner GWAS database (http://phenoscanner.medschl.cam.ac.uk) [19,20] to assess any previous associations (*P* < 0.0033 (0.05/15)) with 4 plausible confounders selected based on previously published studies: birth weight [15,21,22], years of educational attainment and age completed full time education [23–25], and maternal smoking around birth [26–29]. Two SNPs were associated with a potential confounder (rs12041852, maternal smoking around birth, *P* = 7.43 × 10^−5^; rs12507026 (in LD with rs13130484), years of educational attainment, *P* = 0.0028), resulting a set of 13 SNPs for further analysis. In addition to the GWAS which reported these SNPs, 11 of the 13 loci have also been reported to be associated with childhood obesity in other previously published studies (S3 Table). For each outcome, we also used the RadialMR [30] package to further exclude outlying pleiotropic SNPs. RadialMR [30] identified outlying genetic instruments via modified Q-statistics. Among the 13 SNPs, 9 SNPs were in LD with adulthood BMI variants. In addition to the analyses with 13 SNPs, we also carried out MR analyses with the rest 4 SNPs to assess the effects of childhood BMI.

### MR analyses

We used four complementary methods of two-sample MR (inverse variance weighted (IVW) method, MR-Egger method, weighted median method, and weighted mode method) to estimate the causal effects. They make different assumptions about horizontal pleiotropy. When the horizontal pleiotropy is balanced (i.e., the pleiotropic effects are independent of SNP-exposure effects), there should be no bias in the effect derived from MR. If the horizontal pleiotropic effects are biasing the estimate in the same direction (directional pleiotropy), the causal estimates will be biased (except for the MR-Egger method).

The IVW method assumes balanced pleiotropy [31]. We obtained the IVW estimate by meta-analyzing the SNP specific Wald estimates using multiplicative random effects. Cochran’s Q statistic [32] was used to check for the presence of heterogeneity, which can indicate pleiotropy. Cochran’s Q statistic[32] follows a χ^2^ distribution with *L* – 1 degrees of freedom (*L* refers to the number of instruments) under the null hypothesis of homogeneity.

The MR-Egger method is based on the INSIDE assumption (instrument strength independent of the direct effects) [31]. It requires that the SNPs’ potential pleiotropic effects are independent of the SNPs’ association with the exposure [31]. MR-Egger is also based on the no measurement error in the SNP exposure effects (NOME) assumption, which can be evaluated by the regression dilution I^2^_(GX)_ [33]. When I^2^_(GX)_ < 0.9, adjustment methods should be considered [33]. Therefore, simulation extrapolation (SIMEX) correction analysis was performed to estimate the causal effect when I^2^_(GX)_ < 0.9 [33]. The intercept term of the MR-Egger method represents an estimate of the directional pleiotropic effect [34]. We also calculated the Rucker’s Q′ statistic [35] to measure the heterogeneity in the MR-Egger analysis. Rucker’s Q′ follows a χ^2^ distribution with *L* – 2 degrees of freedom under the null hypothesis of no heterogeneity (*L* refers to the number of instruments) [35]. Generally, we have Rucker’s Q′ ≤ Cochran’s Q [35]. If the difference Q – Q′ is sufficiently extreme with respect to a χ^2^ distribution with the 1 degree of freedom, we would infer that directional pleiotropy is an important factor and MR-Egger model provides a better fit than the IVW method [36].

The weighted median method estimates the causal effect under the assumption that at least 50% of the total weight of the instrument comes from valid variants [37]. Compared with IVW and MR-Egger, this method has greater robustness to provide a consistent causal effect estimate even when up to 50% of the SNPs are invalid instruments [37]. The mode-based method provides a consistent effect estimate when the largest number of similar individual-instrument estimates come from valid instruments, even if the majority of instruments are invalid [38].

We also used MR pleiotropy residual sum and outlier (MR-PRESSO) global test [39] to detect horizontal pleiotropy. The analyses of the four MR methods were carried out using the TwoSampleMR package in R. We chose the main MR method as follows:

a. If no directional pleiotropy was detected (*P* > 0.05 for tests of Q, MR-Egger intercept, Q – Q′ and MR-PRESSO), use IVW.
b. If directional pleiotropy was detected and *P* > 0.05 for the test of Q′, use MR-Egger.
c. If directional pleiotropy was detected and *P* < 0.05 for the test of Q′, use weighted median. Effect estimates are reported in β values for continuous outcomes and converted to ORs for dichotomous outcomes.

### Sensitivity analysis

For outcomes with significant MR analysis results, leave-one-out sensitivity analysis was carried out to check whether the causal association was driven by a single SNP.

### Estimate the number of independent outcomes

As our analysis involved a large number of summary data, we expected that some of these outcomes might be highly correlated with each other. Therefore, we used PhenoSpD [40] to estimate the number of independent outcomes to correct for multiple testing. We used the LD score regression method [18] to create a correlation matrix between each outcome. The matrix was used as an input for PhenoSpD to assess the number of independent outcomes through matrix spectral decomposition. Suppose the number of independent outcomes is *n*, then the significant threshold was set as 0.05/*n* after multiple testing correction.

### Instruments for adulthood BMI

For outcomes with significant MR analysis results, we also carried out MR analyses for adulthood BMI. Seventy-six independent SNPs identified by GWAS in populations of European ancestry for adulthood BMI [41] were used as instruments (S4 Table). To investigate whether the relationships between adulthood BMI and other traits/diseases were independent of childhood obesity, we re-ran the MR analysis excluding 11 SNPs included in or in LD (r^2^ > 0.8) with the childhood BMI SNPs (S4 Table).

### Network MR analyses

Considering that lifestyle factors (dietary habits, physical activities, and smoking/drinking behaviors) are also associated with increased CAD/T2D risk, we investigated whether the relationships between childhood BMI and CAD/T2D were mediated by altered lifestyle factors. The summary dataset for CAD [42] was from a meta-analysis of GWAS studies of mainly Europeans, including 60,801 CAD cases and 123,504 controls. The summary data for T2D was from a study [43] combined data from 74,124 T2D cases and 824,006 controls of European ancestry. Lifestyle factors that were significantly affected by childhood BMI in the MR analyses were chosen as potential mediators. The summary data for lifestyle factors were derived from the UK-biobank. MR analyses using lifestyle factors as exposures and CAD/T2D as outcomes were firstly performed. Bi-directional MR were performed to rule out potential reverse causal effects. For each exposure, we used the clumping algorithm in PLINK [44] to select independent SNPs for each trait (r^2^ threshold = 0.001, window size = 1 Mb and *P* < 5 × 10^−8^). The 1000G European data (phase 3) were used as the reference for LD estimation. For exposures with less than 3 significant SNPs available for MR, we used SNPs meeting a more relaxed threshold (*P* < 1 × 10^−5^). This relaxing statistical threshold method for genetic instruments has been used in previous MR studies [45]. The MR analyses process was the same as previously described.

Next, we used the following regression-based method formula [46] for mediation analyses to quantify how much of the association between childhood BMI and CAD/T2D is mediated by selected lifestyle factors:

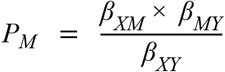

Where *P_M_* represents the proportion of the mediation effect, *β_XM_* refers to the effect of X (childhood BMI) on the mediator (lifestyle factors), *β_MY_* refers to the effect of mediator on CAD/T2D, and *β_XY_* refers to the effect of childhood BMI on CAD/T2D. All effect values were obtained from MR analyses.

## Results

### Genetic correlation analyses

According to the cross-trait LD score analyses, 269 outcomes showed genetic correlation with childhood BMI (Fig 1B, S5 Table), including 254 outcomes from the UK Biobank population. We manually checked the cohorts involved in these outcomes and found that samples in these studies were not overlapped with those in the childhood BMI study. These outcomes (134 disease-related traits, 80 lifestyle factors, and 55 other traits) were subjected to subsequent MR analysis.

### Assessment of pleiotropy

The results of assessment of pleiotropy are shown in S6 Table. No significant evidence of pleiotropy was detected by the Cochran’s Q test and MR-PRESSO global test (*P* > 0.05). MR-Egger’s intercept test detected evidence of directional pleiotropy for 3 outcomes (*P* < 0.05, S5 Table, S1A Fig). The difference Q – Q′ is sufficiently extreme with respect to a χ^2^ distribution with the 1 degree of freedom in additional 7 outcomes (*P* < 0.05, S6 Table, S1B Fig). Since Rucker’s Q′ test didn’t detect evidence of heterogeneity in these 10 outcomes, MR-Egger was chosen as the main method for them. For the rest 259 outcomes without evidence of directional pleiotropy, we chose IVW as the main MR method.

The NOME assumption violation (I^2^_(GX)_ < 0.9) was detected in all outcomes (S5 Table). Therefore, we also carried out MR-Egger with SIMEX analyses.

### MR results

The results of PhenoSpD showed that the independent outcome number was 148, setting the Bonferroni *P*-value threshold for our main MR analysis at *P* < 3.38 × 10^−4^ (0.05/148). In addition to multiple testing corrections of the main MR method, *P* < 3.38 × 10^−4^ of the weighted median method was also set as a cutoff to obtain confident results supported by at least two MR methods. 60 significant associations were detected (S7 Table). Among these 60 traits, 3 traits (CAD, HDL cholesterol and T2D) were from other data resources rather than the UK Biobank population. A total of 27 disease-related traits, 27 lifestyle factors, and 6 other traits were included. For better illustration, we summarized the significant MR findings in Fig 2.

**Fig 2.**
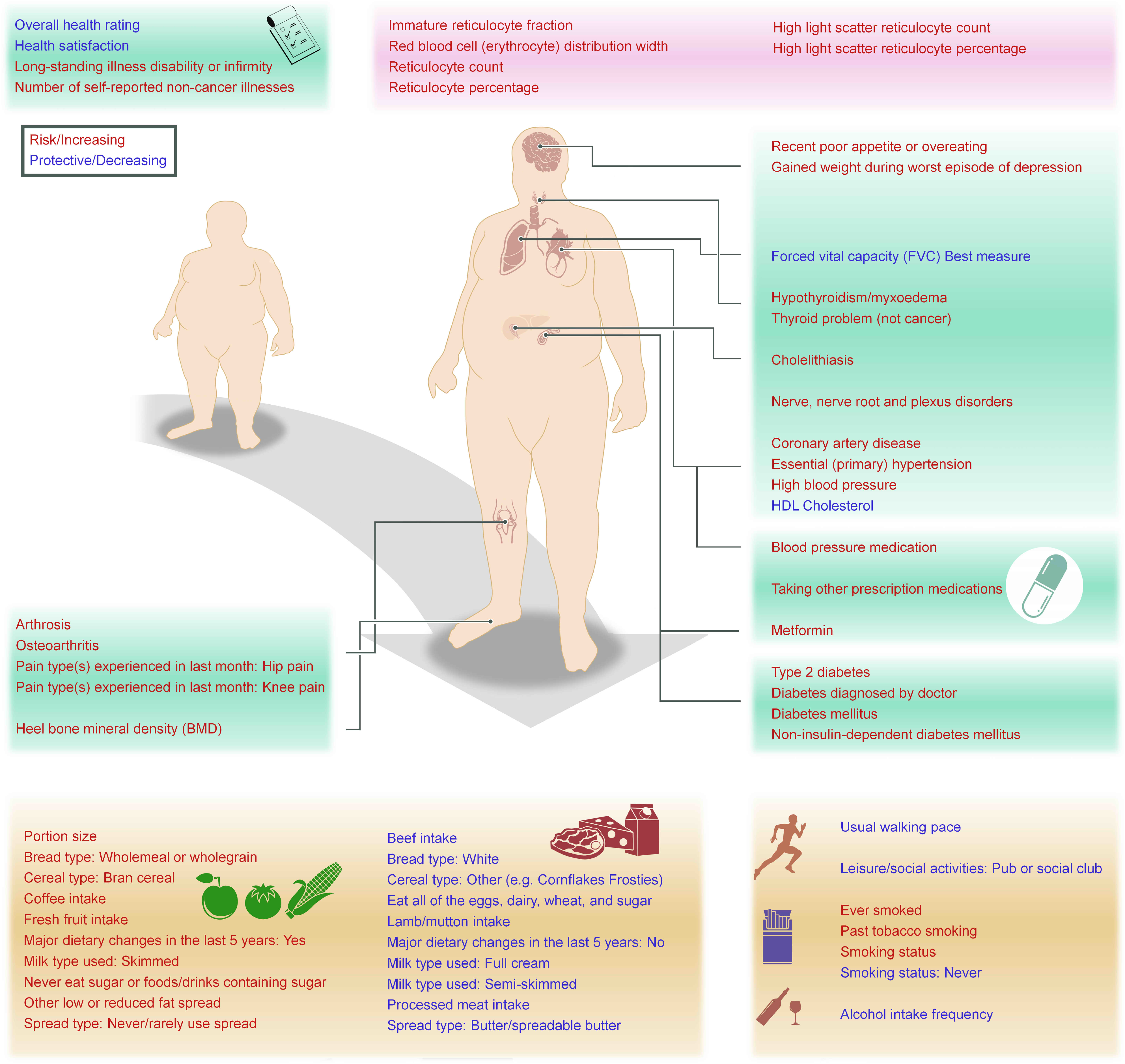
Summary view in which our analyses have shown significant evidence of a causal link between higher childhood BMI and adulthood outcomes. The background colors for disease related traits, lifestyle factors and others are shown in green, yellow and pink, respectively.

The performances of the four methods were similar. Using the threshold of *P* < 3.38 × 10^−4^, the IVW and weighted median methods supported the causal associations between childhood BMI and all 60 traits. But the numbers of associations supported by the weighted mode and MR-Egger methods were only 1 and 3 outcomes, respectively. The difference may be due to the fact that the power of weighted mode and MR-Egger methods is smaller than that of the IVW and weighted median methods [38]. At the suggestive significant level of 0.05, 59 of the 60 associations were supported by at least three methods. The weighted mode and MR-Egger method detected the associations with 58 and 29 outcomes, respectively. This is consistent with the previous report that MR-Egger has the lowest power of the four methods to detect a causal effect [38].

#### Childhood obesity is a risk factor for general health outcomes and socioeconomic status in adulthood

As shown in Fig 3, higher childhood BMI was associated with reduced overall health rating (β = −0.10, 95% CI: −0.13 to −0.07, *P* = 6.26 × 10^−11^), health satisfaction (β = −0.13, 95% CI: −0.18 to −0.08, *P* = 7.44 × 10^−7^) and an increased the number of self-reported non-cancer illnesses (β = 0.09, 95% CI: 0.06 to 0.13, *P* = 1.58 × 10^−7^). One SD increase in childhood BMI was associated with 9% higher odds of long-standing illness disability or infirmity (OR = 1.09, 95% CI: 1.06 to 1.12, *P* = 8.50 × 10^−11^). Higher childhood BMI has a causal relationship with lower average total household income before tax (main MR method β = −0.06, 95% CI: −0.09 to −0.03, *P* = 8.54 × 10^−5^), although the weighted median test for this trait was not significant after multiple testing correction (*P* = 9.46 × 10^−4^). Leave-one-out analysis showed that no single SNP was driving the causal estimates (S2 Fig). Therefore, childhood obesity is a risk factor for general health outcomes and socioeconomic status.

**Fig 3.**
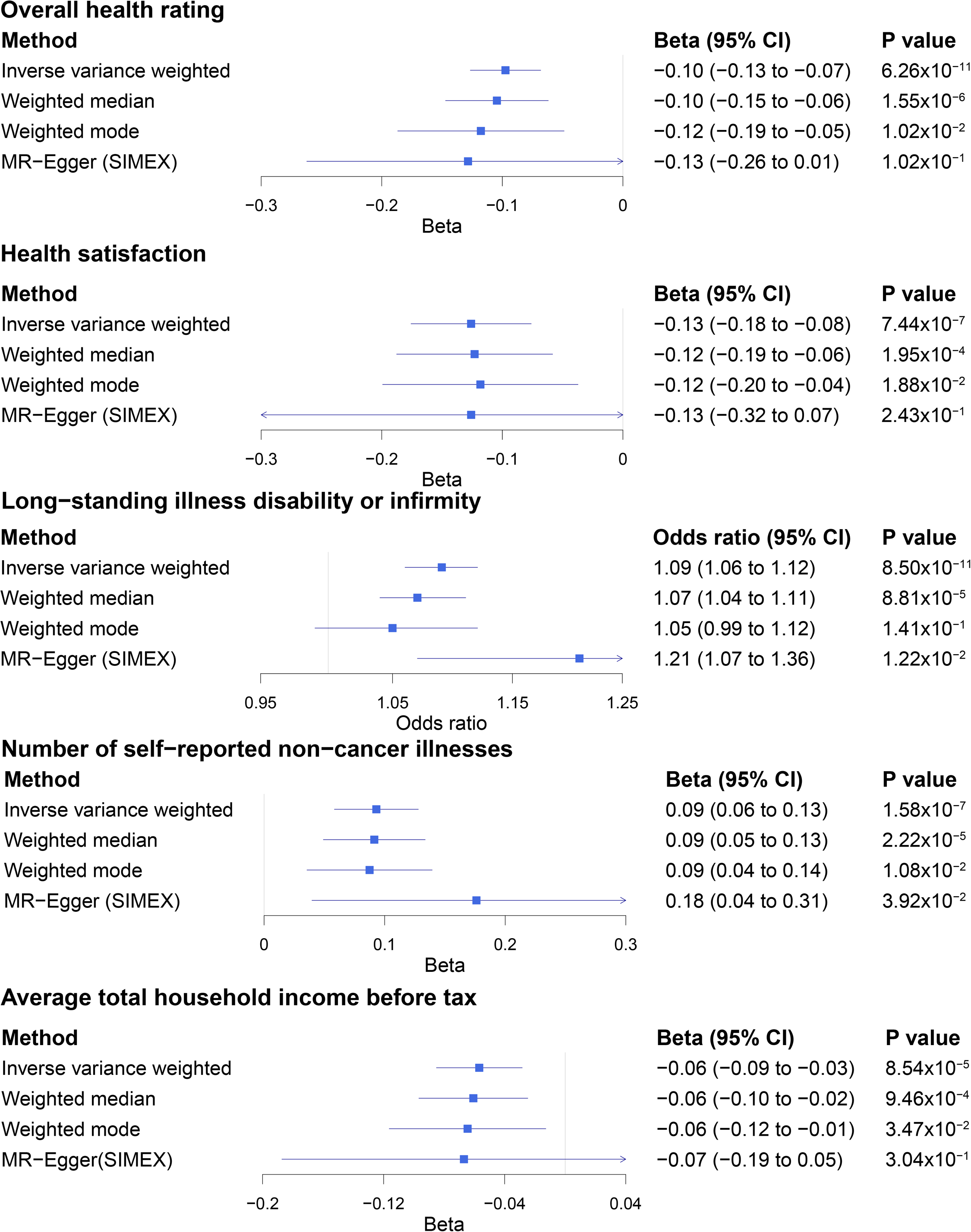
Summary Mendelian randomization (MR) estimates derived from the main inverse-variance weighted, MR-Egger, weighted median and weighted mode-based methods for overall heath rating, health satisfaction, long-standing illness disability or infirmity, number of self-reported non-caner illnesses and average total household income before tax.

##### Childhood BMI and adult diseases-related traits

There is evidence that childhood BMI causally affects a total of 27 outcomes related to adult diseases, including 3 circulatory system traits; 7 endocrine, nutritional, metabolic traits; 5 musculoskeletal system traits; and 10 other traits.

As shown in Fig 4 and S3 Fig, we found evidence that higher childhood BMI caused an increased risk of cholelithiasis (OR = 1.26, 95% CI: 1.18 to 1.35, *P* = 3.29 × 10^−5^), and the risk effect was supported by three MR methods (IVW, weighted median and MR-Egger) after multiple testing corrections. We also observed adverse effects of childhood BMI on hypothyroidism (OR = 1.06, 95% CI: 1.03 to 1.09, *P* = 8.77 × 10^−6^) and non-cancer thyroid problems (OR = 1.07, 95% CI: 1.04 to 1.10, *P* = 7.78 × 10^−7^). We also found evidence that childhood BMI was positively associated with adult heel bone mineral density (BMD) (β = 0.20, 95% CI: 0.15 to 0.24, *P* = 3.40 × 10^−20^). Our results help to clarify the association between childhood BMI and later-life osteoporosis.

**Fig 4.**
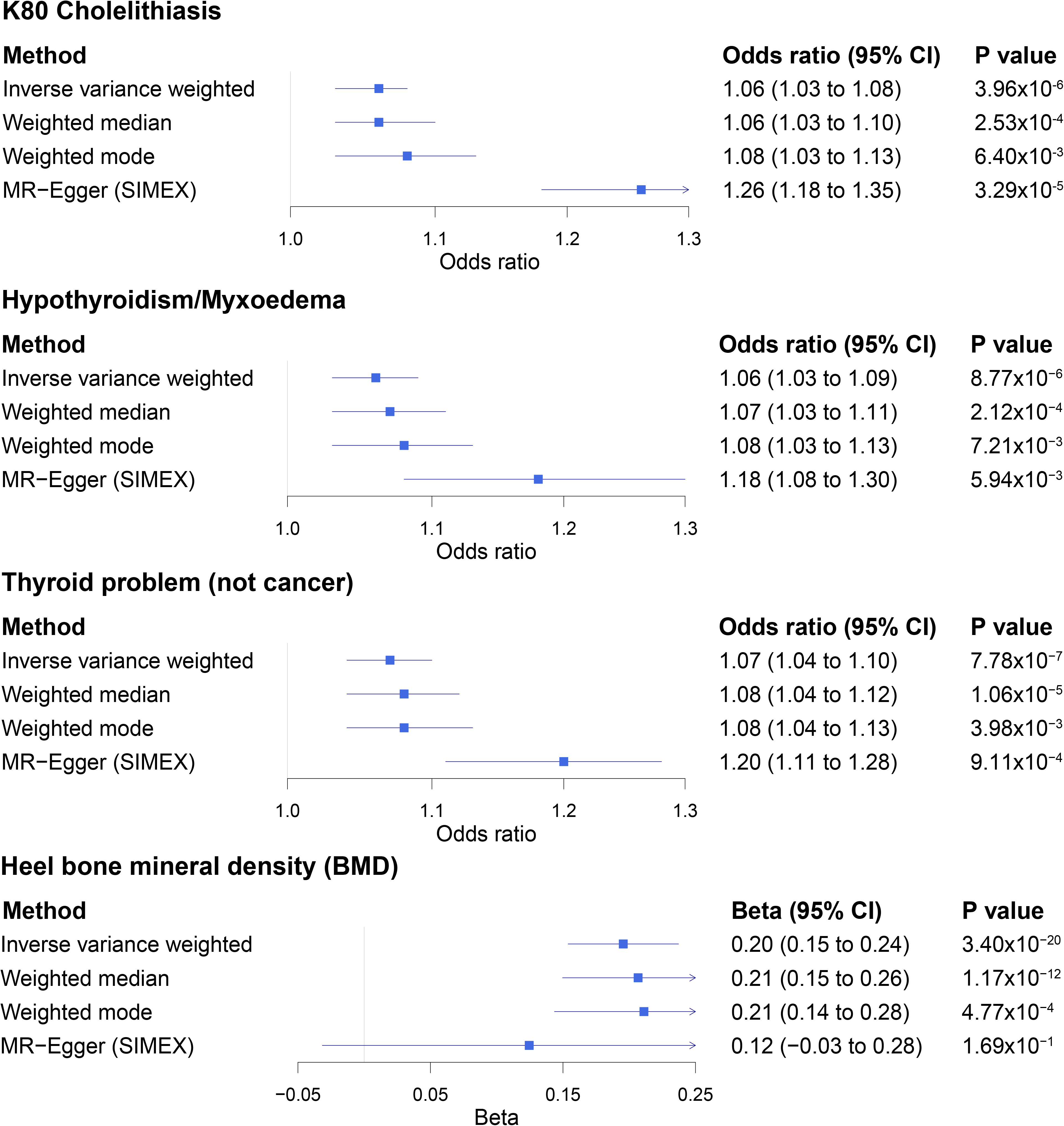
Summary Mendelian randomization (MR) estimates derived from the main inverse-variance weighted, MR-Egger, weighted median and weighted mode-based methods for cholelithiasis, hypothyroidism, thyroid problem (not cancer), and heel bone mineral density.

Consistent with previous findings, we found that a 1 SD increase in childhood BMI was associated with 10% higher odds of CAD (OR = 1.10, 95% CI: 1.06 to 1.15, *P* = 1.20 × 10^−6^, S4 Fig). The other two circulatory system traits (essential hypertension and high blood pressure) are all closely related with CAD (S4 Fig). Analyses of treatment/medication conditions also showed that higher childhood BMI increased the risk of receiving blood pressure medication (S5 Fig). Consistent with previous findings, we also observed that a 1 SD increase in childhood BMI was associated with 36% higher odds of T2D (OR = 1.36, 95% CI: 1.30 to 1.43, *P* = 1.57 × 10^−34^, S6 Fig). We found evidence that higher childhood BMI caused increased risk of T2D-related traits (S6 Fig). Higher childhood BMI also increased the risk of receiving Metformin, the drug for T2D treatment (S5 Fig). The risk effects on high blood pressure and T2D were supported by three MR methods after multiple testing corrections. In addition, we confirmed the known risk effects of childhood BMI on osteoarthritis (S7 Fig).

#### Childhood BMI and adult lifestyle factors

There is evidence that childhood BMI causally affects a total of 27 adult lifestyle factors, including 20 dietary habits, 4 smoking behaviors, usual walking pace, pub/social club activities, and alcohol intake frequency.

##### Childhood BMI and adult dietary habits

As it might be expected, we observed a positive association between childhood BMI and adult diet portion size (β = 0.26, 95% CI: 0.18 to 0.34, *P* = 7.34 × 10^−11^, S7 Table, S8 Fig). However, it might be unexpected that higher childhood BMI was positively associated with low calorie density food intake (S7 Table, S8 Fig). For example, childhood BMI was positively associated with the intake of high-fiber foods (e.g., fresh fruit intake, bran cereal, and wholemeal bread) and low fat/sugar food (e.g., skimmed milk, never/rarely using spread on bread, never eat sugar or food/drinks containing sugar). We also found negative associations between childhood BMI and the intake of meat (beef, lamb/mutton, and processed meat), full cream milk, and butter spread on bread.

##### Childhood BMI and adult physical activities, smoking/drinking behaviors

As shown in Fig 5 and S9 Fig, for physical activities, we noticed that childhood BMI was negatively associated with usual walking pace (β = −0.12, 95% CI: −0.15 to −0.08, *P* = 3.24 × 10^−10^). For smoking behaviors, we observed positive associations between childhood BMI and adult smoking status. In contrast, higher childhood BMI was negatively associated with alcohol intake frequency (β = −0.13, CI: −0.17 to −0.09, *P* = 2.74 × 10^−11^).

**Fig 5.**
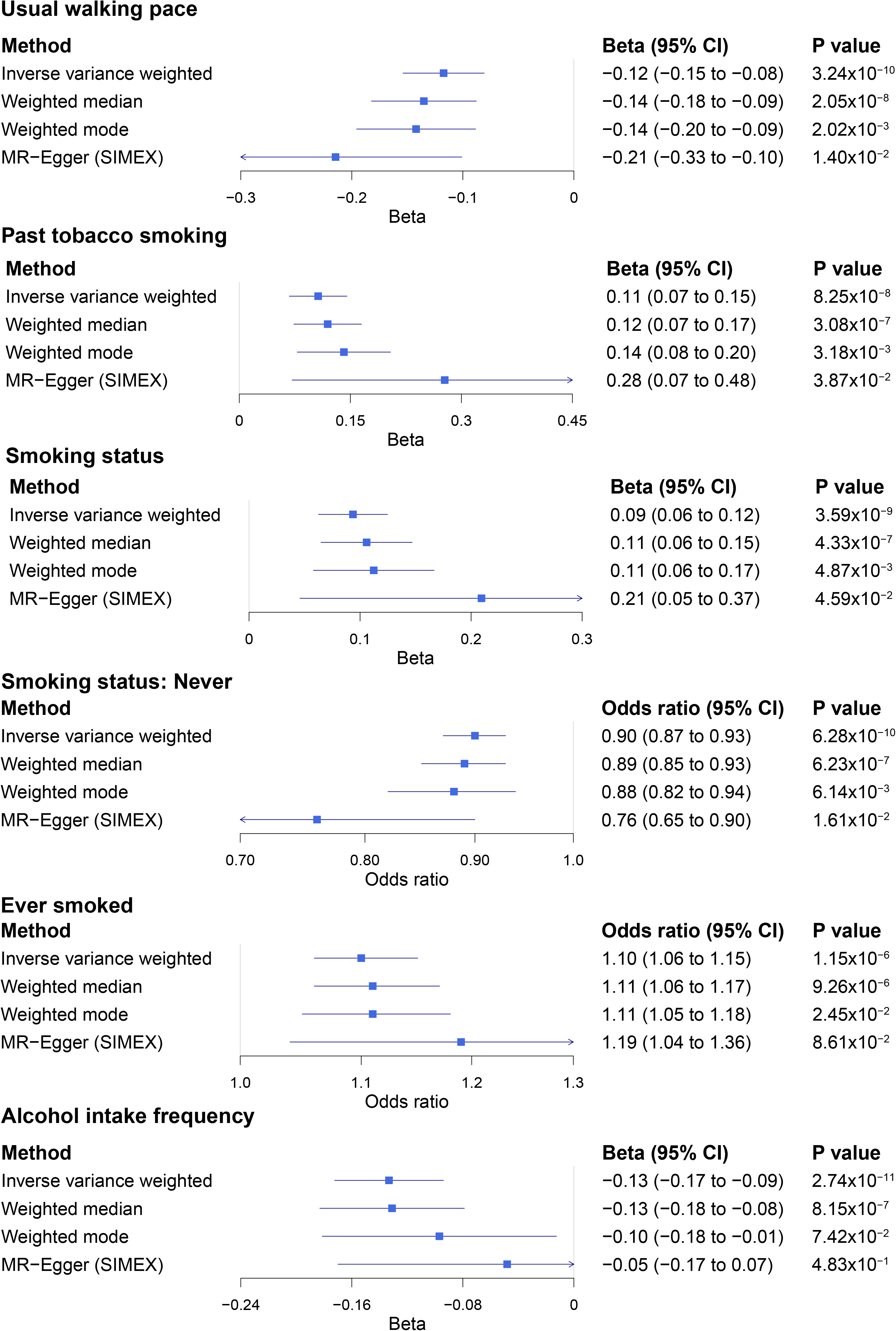
Summary Mendelian randomization (MR) estimates derived from the main inverse-variance weighted, MR-Egger, weighted median and weighted mode-based methods for usual walking pace, past tobacco smoking, smoking status, smoking status: never, ever smoked, and alcohol intake frequency.

#### MR analysis for childhood BMI after removing loci associated with adult BMI

For the 60 outcomes with significant MR analysis results, we also performed MR analyses using the 4 SNPs which were not in LD with the adult BMI variants, to assess the causal effects of childhood BMI which might be independent of adult BMI. As shown in S8 Table, due to the limited number of SNPs, the effects on adult traits were attenuated or no longer present. At the significant level of *P* < 0.05, we detected the associations with 13 traits (21.67%, 13/60). These outcomes included 4 T2D related traits, 7 dietary habits, 1 trait about general health and 1 blood count related trait. For example, higher childhood BMI increased the risk of receiving Metformin (drug for T2D treatment) (OR = 1.11, 95% CI: 1.04 to 1.18, *P* = 7.13 × 10^−4^). Negative association between childhood BMI and butter spread on bread was observed (OR = 0.91, 95% CI: 0.86 to 0.96, *P* = 1.50 × 10^−3^).

#### Positive association between adulthood BMI and heel BMD no longer present after removing loci associated with childhood BMI

For the 60 outcomes with significant MR analysis results, we also carried out MR analyses for adulthood BMI. As it might be expected, although the effect sizes were different, at least suggestive associations (*P* < 0.05) were detected between adulthood BMI and these traits (S9 Table). The results were similar to the results of Millard *et al*.[12] We next analyzed whether the effects of adulthood BMI are independent of childhood BMI. After excluding SNPs that existed in or in LD with childhood BMI, the associations between adulthood BMI and 3 traits vanished (*P* > 0.05 in all MR analyses method, S10 Table), suggesting that the causal associations between these 3 traits and adulthood BMI might depend on childhood obesity. For example, with the original 76 SNPs as instruments, adulthood BMI was positively associated with heel BMD (β = 0.10, CI: 0.05 to 0.15, *P* = 5.31 × 10^−5^). However, the association no longer present after excluding the 11 SNPs that existed in or in LD with childhood BMI (*P* > 0.05).

To test whether the results of removing 11 childhood BMI SNPs might be underpowered and biased towards SNPs which have small/marginal effects on BMI, we carried out the following analysis. We randomly selected 1,000 sets of 11 SNPs with similar β values comparing to the 11 childhood BMI SNPs (Wilcox test *P* > 0.05) from the rest 65 SNPs. Then we performed 1,000 MR analyses after removing these random sets from the original 76 SNPs for the 3 outcomes whose associations were no long present. For heel BMD, all of the 1,000 MR analyses remained significant at *P* < 0.01 level, and the *P* values of 998 tests were less than 1 × 10^−4^ (median *P* = 3.86 × 10^−8^). Therefore, it is likely that the 11 childhood BMI SNPs contribute most to the detected protective effect on heel BMD.

#### Childhood BMI, adult lifestyle factors, and CAD/T2D: network MR analyses

As shown in Fig 6A, among the 27 lifestyle factors, genetic correlation analyses showed that 20 and 22 were genetically correlated with CAD and T2D, respectively (S11 Table). The results of PhenoSpD showed that the independent trait numbers were 13 and 14 respectively, setting the significant threshold for network MR analyses as *P* < 9.26 × 10^−4^ (0.05 / ((13 + 14) × 2)). Lifestyle factors or socioeconomic status with significant effects on CAD/T2D and without reverse causal effects were selected as potential mediators.

**Fig 6.**
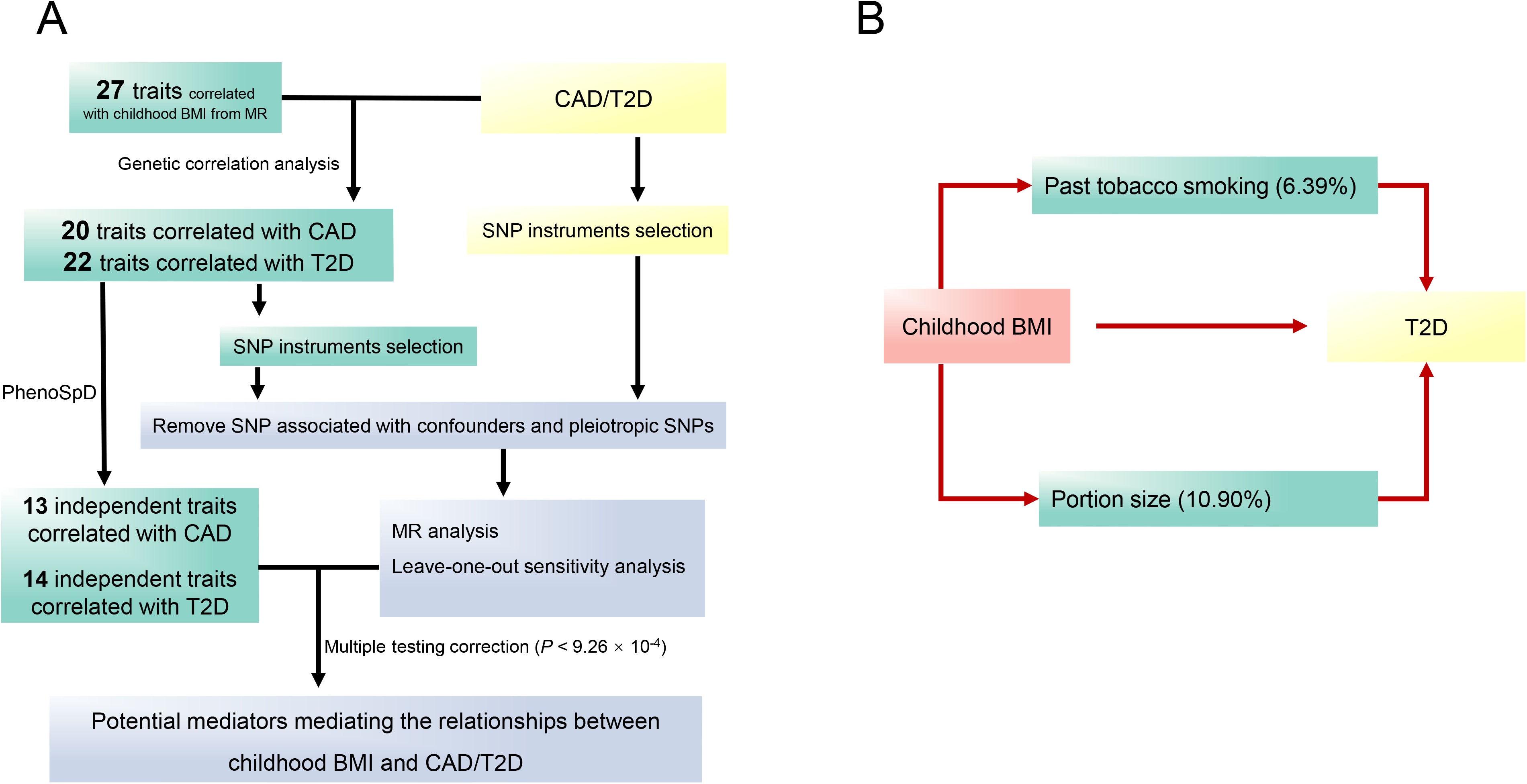
A. The network Mendelian randomization (MR) analyses pipeline. B. Mediation effect (proportions shown in brackets) of past tobacco smoking and portion size on the childhood BMI and T2D associations. Past tobacco smoking is a categorical trait which was recoded from 1 to 4 to refer “I have never smoked”, “just tried once or twice”, “smoked occasionally”, and “smoked on most or all days”, respectively.

After multiple testing corrections, none traits showed a causal association with CAD. Therefore, we did not find suitable mediators for CAD. For T2D, as shown in S12 Table, 7 lifestyle factors showed significant causal effects on T2D. Among them, there were no reverse effects of T2D on past tobacco smoking and portion size (S13 Table, *P* > 0.05 in all MR methods). They were estimated to mediate 6.39% and 10.90% of the associations between childhood BMI and T2D, respectively (Fig 6B).

## Discussion

In this study, with GWAS summary data from public resources, we carried out 2-sample MR analyses to investigate the causal effects of childhood BMI on adult outcomes with genetic correlation. We identified potential causal effects of childhood obesity on 60 adult traits. Compared with previous studies of childhood BMI which only focused on a few traits [10,11], here we provided a phenome-wide investigation of the causal associations between childhood BMI and adult outcomes.

### Childhood obesity is a risk factor for general health outcomes in adulthood

We observed that childhood obesity is a risk factor for general health outcomes in adulthood. Consistently, previous studies have demonstrated that high childhood BMI was associated with increased mortality and morbidity [2] in adulthood.

Specifically, we observed adverse effects of higher childhood BMI on CAD and T2D. This is consistent with the results of a previous MR study by Geng *et al*. [10]. We also replicated their finding about the negative association between childhood BMI and HDL cholesterol level, which is a well-known trait inversely related with CAD [47]. In addition, positive association between childhood BMI and high blood pressure was supported by different MR methods. Our analyses on treatment/medication conditions further showed that higher childhood BMI increased the risk of receiving CAD and T2D related medications, including blood pressure medication and metformin. Observational studies have also shown that higher childhood BMI is related to increased incidence of diabetes [48], CAD [4], and hypertension [49]. These data supported that childhood obesity might be a determinant of adult CAD/T2D risk.

Consistent with another MR study on childhood BMI [11], we detected positive association between childhood BMI and adult osteoarthritis, especially hip and knee pain. A previous observational study suggested that obesity from childhood had an accumulative effect on knee OA development [50]. Similarly, a study by McFarlane *et al*. [51] on the 1958 British birth cohort observed a significant association with knee pain at the age of 45 years with high BMI from as early as age 11 years [51]. Moreover, another study [52] reported that the childhood overweight measures were significantly associated with adulthood knee mechanical joint pain among males, and this association was independent of the adult overweight. Therefore, it is possible that the effect of childhood obesity on the knee joint can persist into adulthood.

In our results, the adverse effects of higher childhood BMI on cholelithiasis was consistently observed by different MR methods. Obesity has been established as an important risk factor for cholelithiasis in both children [53] and adults [54]. Previous MR studies [13,55] in adults suggested that elevated BMI was a causal risk factor for symptomatic gallstone disease. However, the causal association between childhood obesity and the risk of cholelithiasis in adult was not reported before. Our result suggested a potential link between childhood obesity and cholelithiasis risk in adults.

### The importance of control portion size is noted

For dietary habits, it was unexpected that childhood obesity is positively associated with low calorie density food in adulthood. However, positive associations between childhood obesity and healthy diet habits have been reported in observational studies previously. For example, a healthy diet score was associated with increased odds of overweight/obesity in children from the UK [56]. Similarly, less frequent intake of energy-dense foods was associated with larger waist circumference in Swedish children [57]. It is possible that subjects suffering from childhood obesity may reduce their intake of unhealthy foods to lose weight, suggesting that advice on improving the quality of eating habits is now reaching the target audience.

It is possible that the identified causal effects of childhood BMI on dietary habits were mediated by other factors, such as socioeconomic status. We did observe that childhood BMI was negatively associated with average total household income before tax. A previous observational study [58] also suggested that childhood obesity adversely affects emotional and social skills, which are important determinants of future economic prosperity. On the other hand, personal income is a key driver of dietary choices. Low-income individuals might face a number of challenges to acquire enough nutritious foods [59]. However, the effect of income on dietary consumption may vary by food category, country, age, and sex [60]. For example, rising income was estimated to increase fruit intake most strongly among older women globally [60]. Current GWAS summary data we used for income and dietary habits are all from the UK Biobank population. Therefore, it is not suitable for two-sample MR analysis, since overlap in participants between the two samples can cause bias towards the risk factor-outcome association [61]. Further data are needed to clarify the potential contribution of socioeconomic status in mediating the association between childhood obesity and adult dietary habits.

In contrast, we observed positive associations between childhood BMI and adult diet portion size. In addition, network MR analyses showed that portion size might mediate the associations between childhood BMI and T2D. Previous observational studies have provided support for the positive association between T2D and binge eating [62]. Our results supported that control portion size may help reduce the risk of T2D.

### Positive association between adulthood BMI and heel BMD no longer present after removing loci associated with childhood BMI

We observed a positive association between childhood BMI and adult heel BMD. A previous MR study reported that adiposity is causally related to increased BMD at all sites except the skull in 5221 subjects from the Avon Longitudinal Study of Parents and Children [63]. In adults, MR analysis suggested that adiposity might be causally related to BMD at the femur [64]. Protective effect on osteoporosis of higher BMI in adults has also been reported previously [65]. We also observed a positive association between adult BMI and adult heel BMD. However, when excluding SNPs existing in or in LD with childhood BMI SNPs, the positive association of adult BMI and heel BMD vanished, suggesting that this association depend on childhood BMI. It is widely accepted that most of the skeletal mass is acquired by the age of 20. Several studies have suggested that peak bone density is achieved by the end of adolescence [66,67]. The risk of developing osteoporosis is influenced to a large extent by the levels of peak BMD. Our results implicated that the increasing effect on BMD of obesity might mainly work in childhood. Investigations taking peak BMD into consideration in adults are further needed to confirm our findings.

### Childhood BMI, adult smoking behavior, and T2D

Network MR analyses showed that past tobacco smoking might mediate 6.39% of the association between childhood BMI and T2D. A variety of epidemiological studies have demonstrated the positive associations between smoking and development of T2D [68,69]. In addition, the risk of diabetes could be reverted in subjects who stopped smoking [70], suggesting that tackling smoking could make contributions to the control of T2D development in adults who once suffered from childhood obesity.

### General limitations of the study

The limitations of the current study should be addressed. Firstly, because there are inevitably overlapping loci between childhood BMI and adulthood BMI, it is hard to identify which of these causal effects are due to early-life obesity, as opposed to late-life effects. Since adulthood BMI is likely affected by childhood BMI and also a cause of many disease outcomes, it is possible that our reported associations could be (partly or exclusively) due to adulthood BMI. However, childhood BMI GWASs conducted to date are notably smaller in sample size compared to adulthood GWASs, it is hard to obtain variants only associated with childhood BMI and not with overall BMI. When data for larger scale GWASs on childhood BMI are available, the power will be improved to identify more SNPs specifically associated with childhood BMI with smaller effects, and then the results of our analysis might be updated. Secondly, although our analyses supported that our results were not affected by pleiotropy, we cannot rule out the possibility of a shared genetic basis rather than a causal relationship. Thirdly, since we used GWAS summary data from the public database for our analyses, we cannot assess the effects of population stratification on our results. Summary data from multiple multi-ethnic populations might lead to biased association results since different ethnic populations have different LD structures and allele frequencies [71]. The summary data we used here were mainly derived from the European population. However, since we did not subset to the European-only results, there is a potential of bias from significant distinctions in disease outcomes between European and non-Europeans. The UK Biobank data were reported to be skewed as wealthier and more educated [72], which might also affect the generalization of the results. Lastly, since large scale sex-specific GWAS on childhood BMI is still lacking, we did not take sex into account in both exposure and outcomes. Besides, our results here primarily assessed the impact of genetically caused changes in childhood BMI and the impact of interventions in later life is likely to be smaller than the estimates here.

## Conclusion

In summary, using public GWAS datasets, we carried out 2-sample MR analyses to investigate the causal effects of childhood BMI on adult outcomes. We identified potential causal effects of childhood obesity on 60 adult traits. Our results highlight the need to intervene in childhood to reduce obesity from a young age and its effects later in life.

## Data sharing

All data is publicly available through data resources indicated in the methods.

## Contributions

T.-L.Y. designed the study. S.-S.D. and Y.G. wrote and edited the manuscript. S.-S.D., K.Z., J.-M.D., J.-C.F., S.Y., F.J., X.-F.C., H.W. and R.-H.H. collected and analyzed the data. J.-M.D., S.Y., J.-B.C. and Y.R. drew the figures.

## Supporting information

**S1 Fig.**
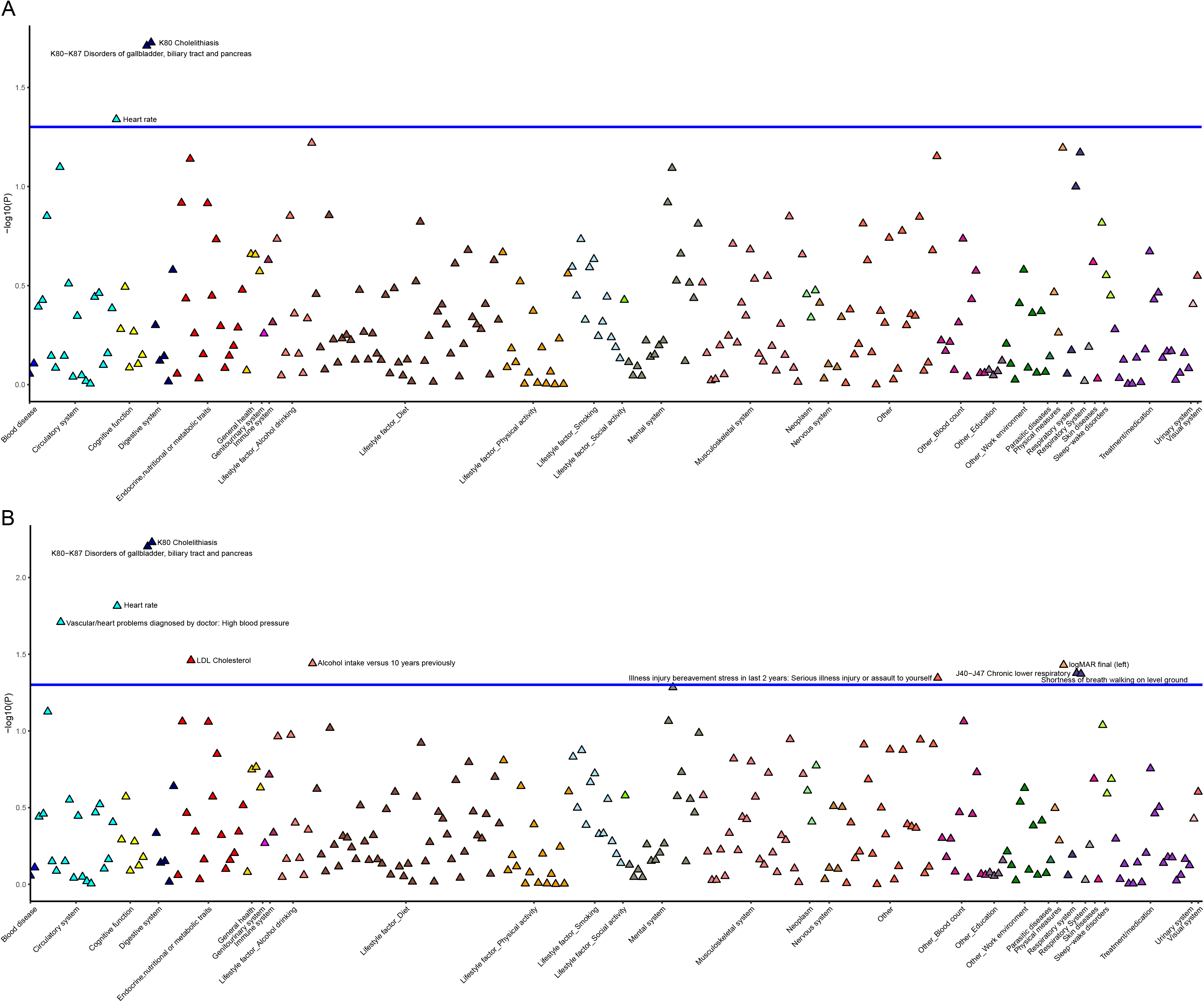
Manhattan plot showing the pleiotropy assessment results for MR-Egger intercept test (A) and Q-Q′ test (B). For each plot, the blue horizontal line represents the threshold for significance (*P* < 0.05).

**S2 Fig.**
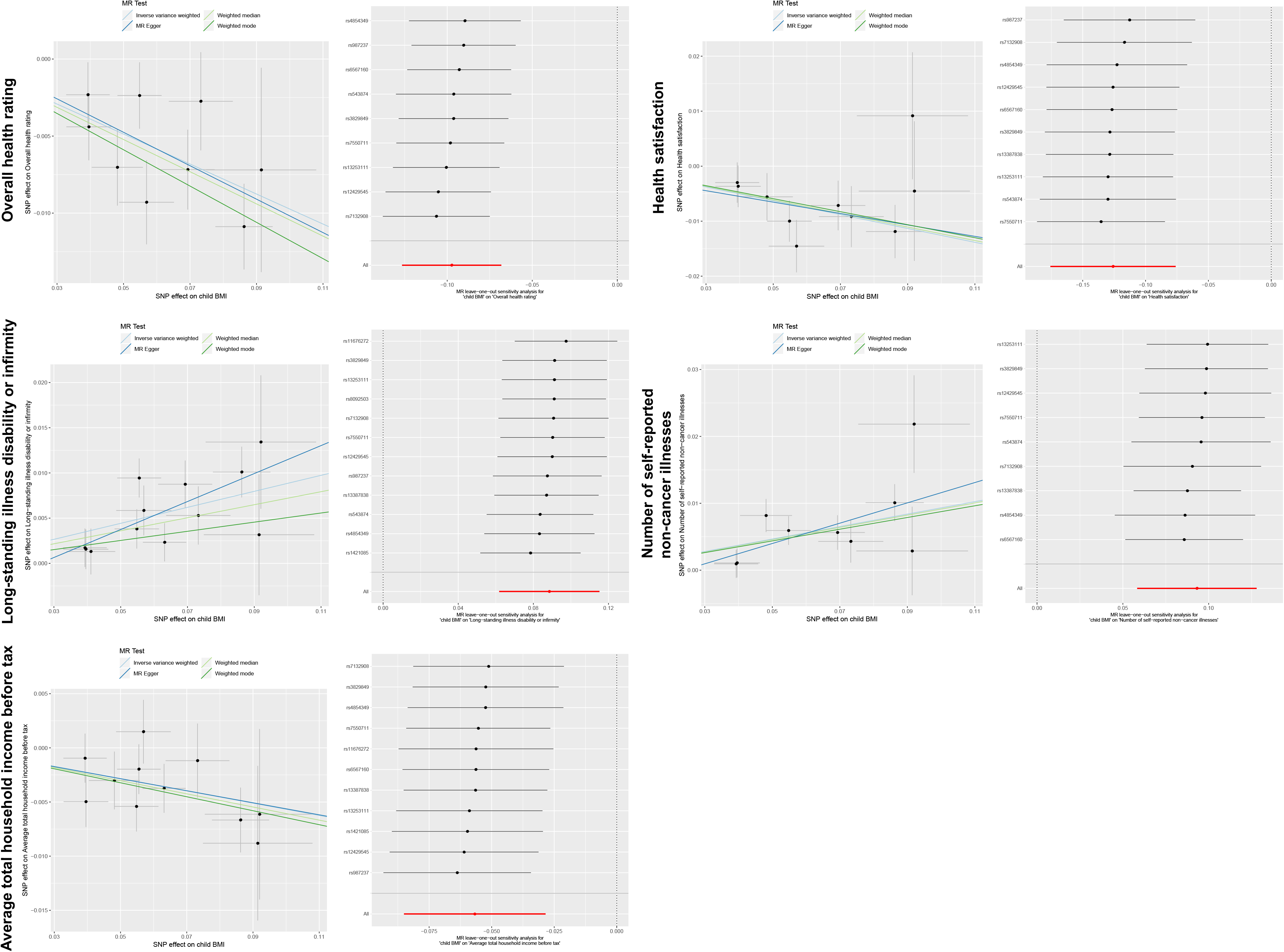
Scatter plot and leave one out analysis plot for overall heath rating, health satisfaction, long-standing illness disability or infirmity, number of self-reported non-cancer illnesses and average total household income before tax.

**S3 Fig.**
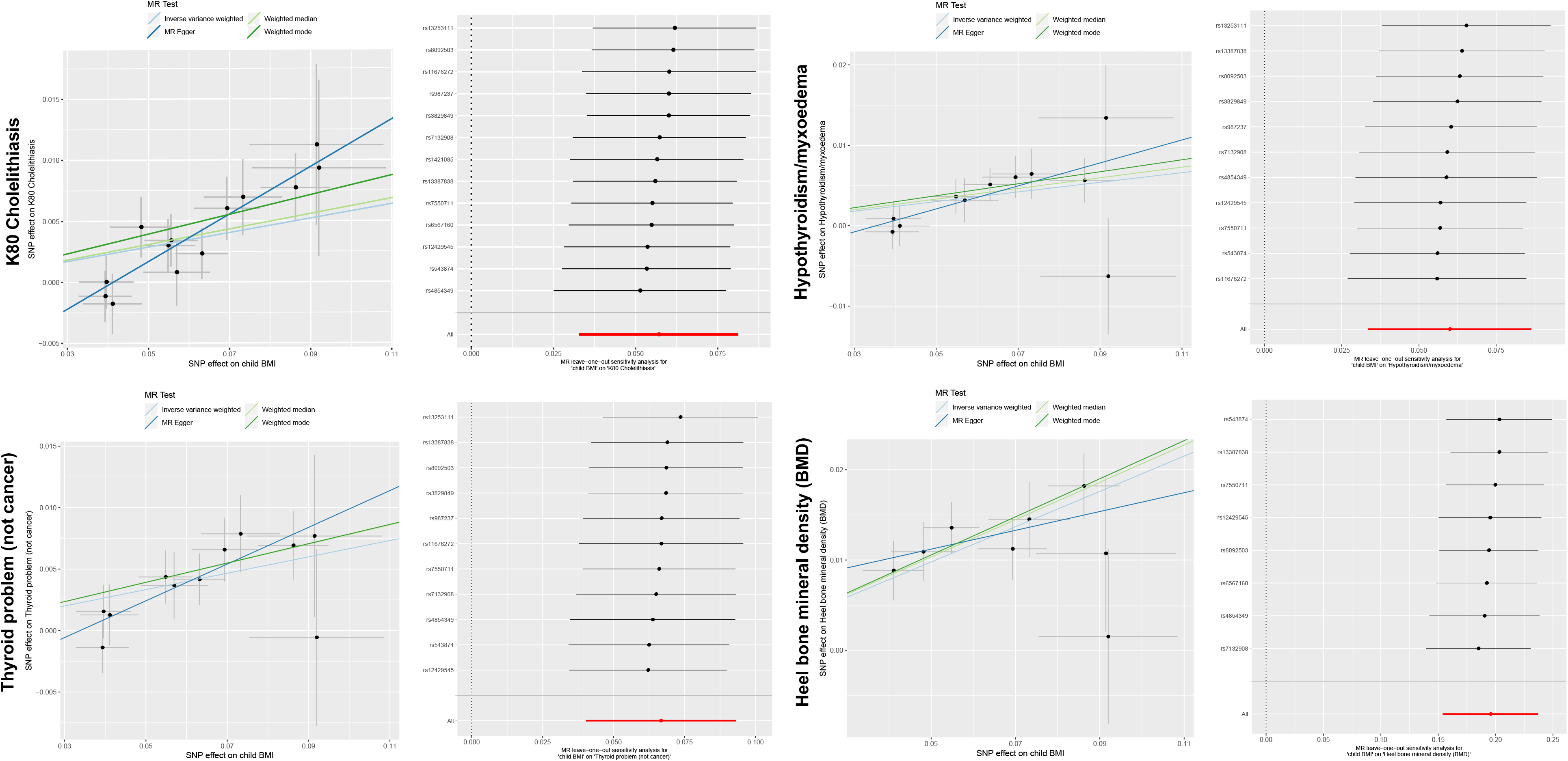
Scatter plot and leave one out analysis plot for cholelithiasis, hypothyroidism, non-cancer thyroid problem and heel bone mineral density (BMD).

**S4 Fig.**
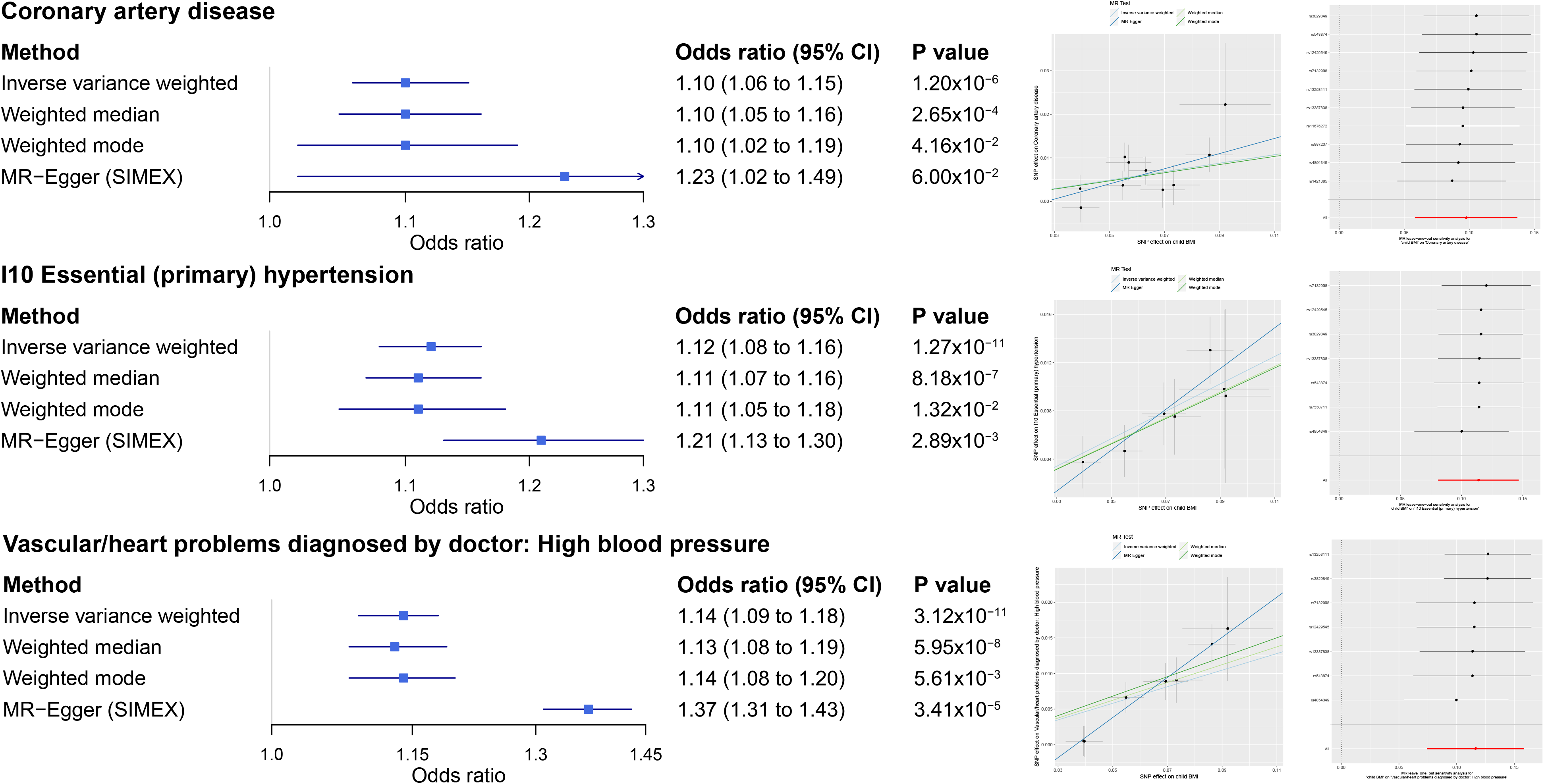
Summary Mendelian randomization (MR) estimates derived from the main inverse-variance weighted, MR-Egger, weighted median and weighted mode-based methods for coronary artery disease and related traits. Scatter plot and leave one out analysis plot for each trait are also shown.

**S5 Fig.**
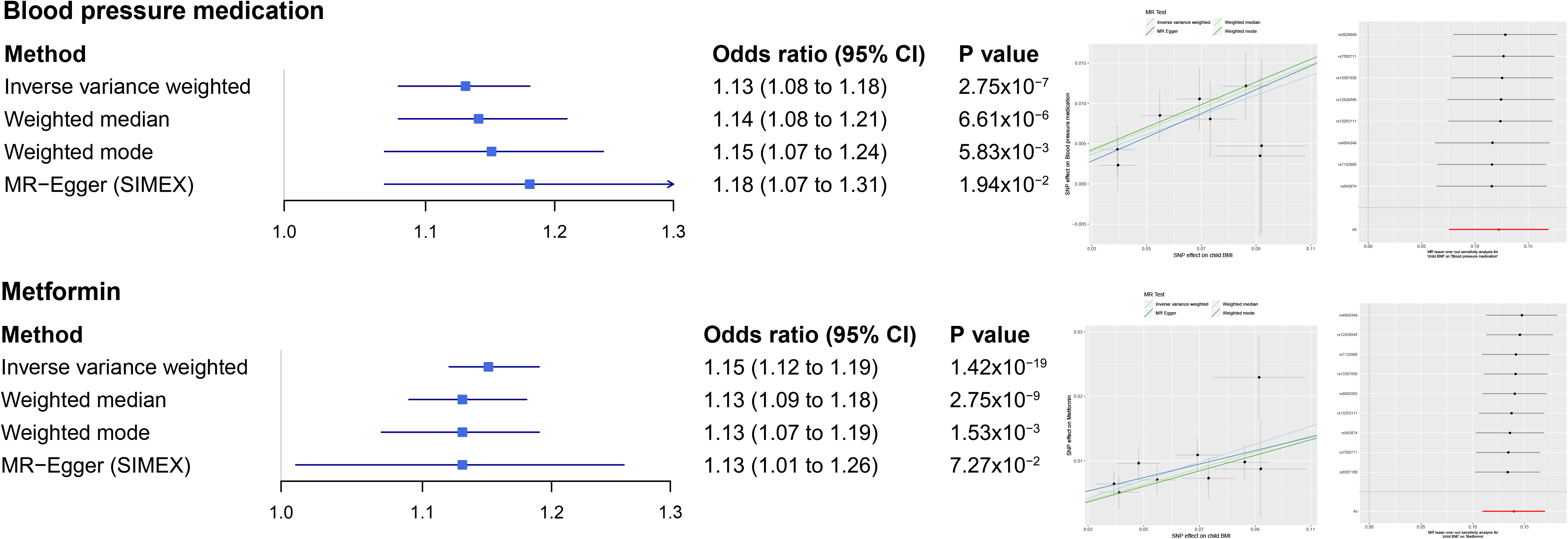
Summary Mendelian randomization (MR) estimates derived from the main inverse-variance weighted, MR-Egger, weighted median and weighted mode-based methods for received treatment/medication. Scatter plot and leave one out analysis plot for each trait are also shown.

**S6 Fig.**
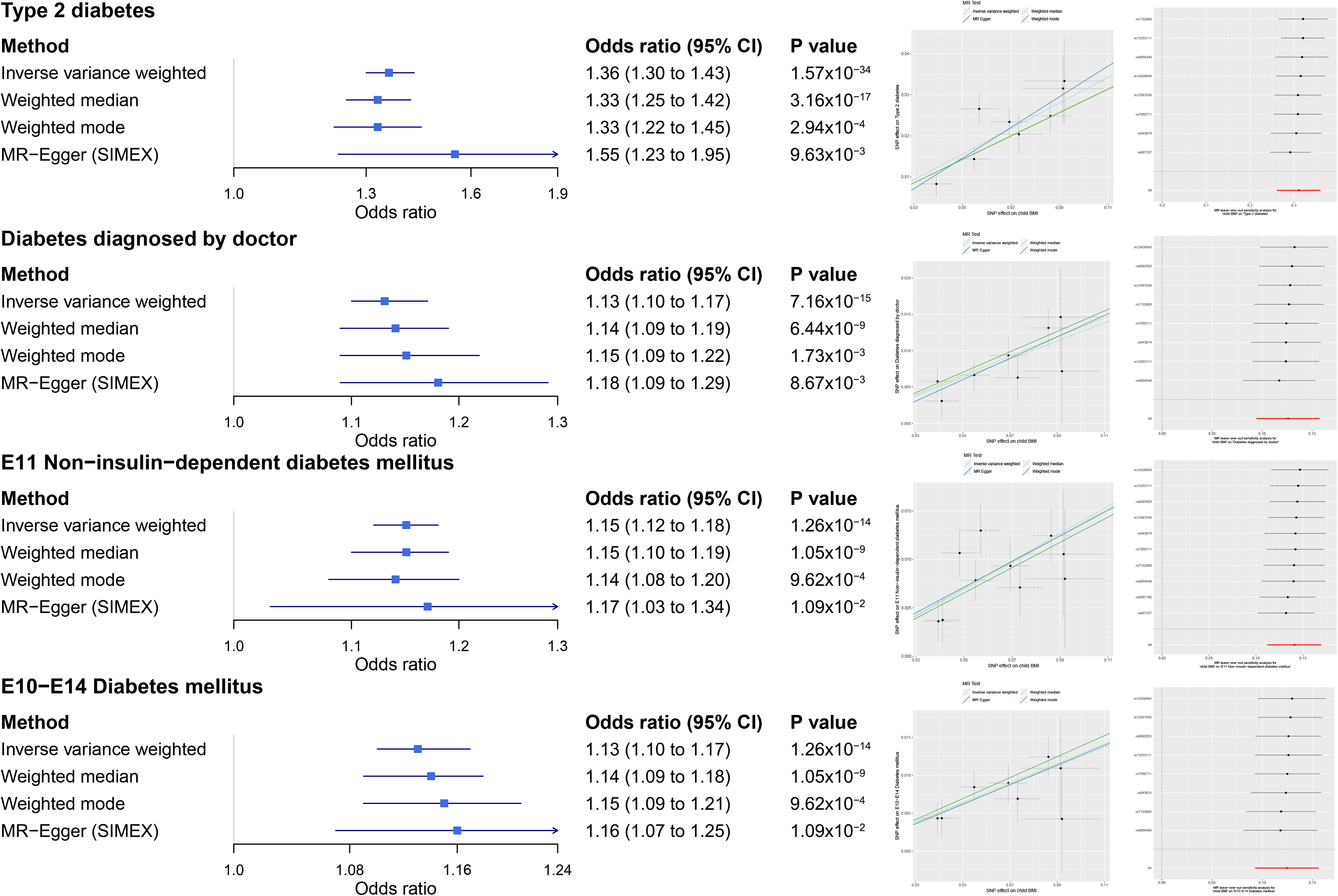
Summary Mendelian randomization (MR) estimates derived from the main inverse-variance weighted, MR-Egger, weighted median and weighted mode-based methods for type 2 diabetes (T2D) and related traits. Scatter plot and leave one out analysis plot for each trait are also shown.

**S7 Fig.**
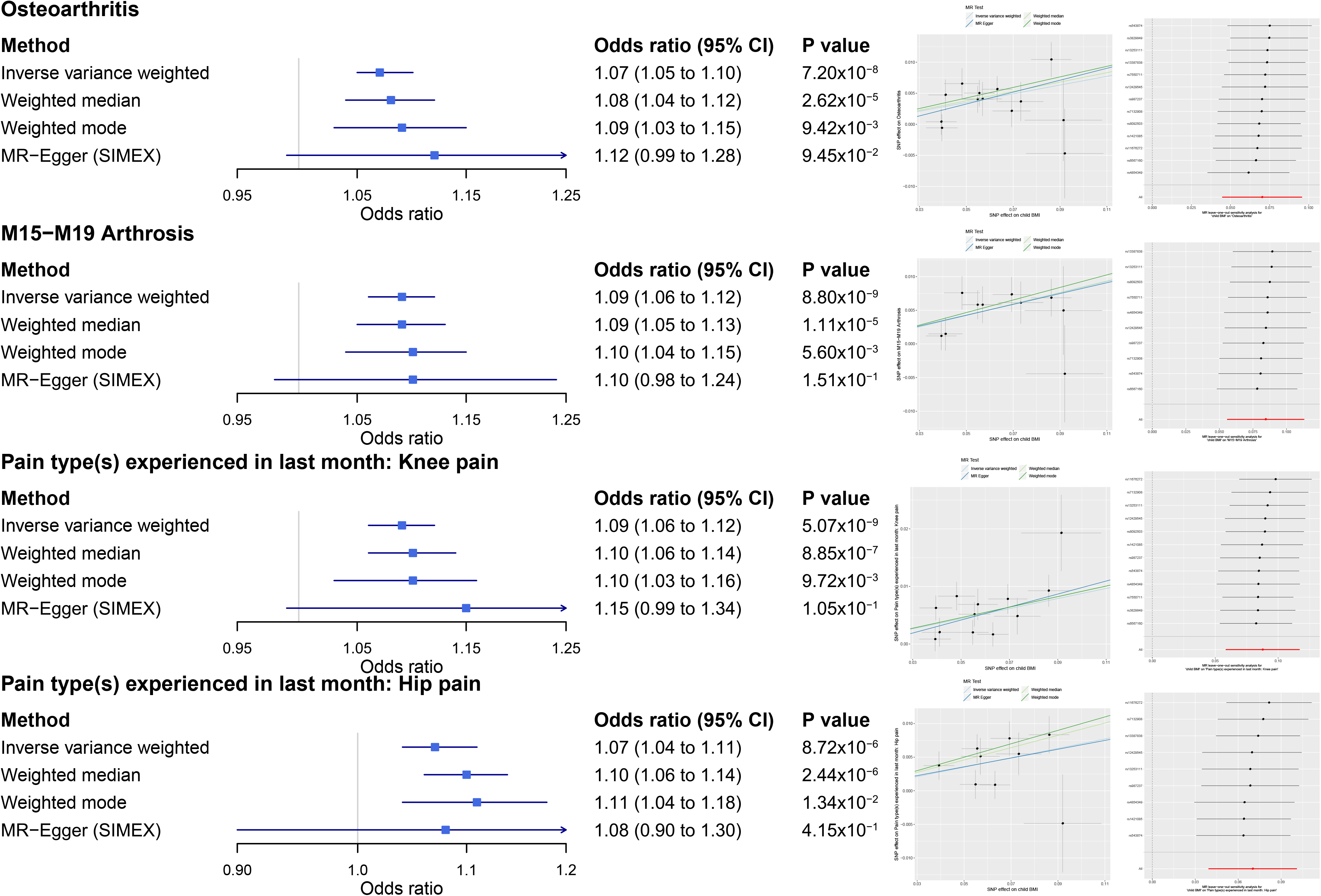
Summary Mendelian randomization (MR) estimates derived from the main inverse-variance weighted, MR-Egger, weighted median and weighted mode-based methods for osteoarthritis and related traits. Scatter plot and leave one out analysis plot for each trait are also shown.

**S8 Fig.**
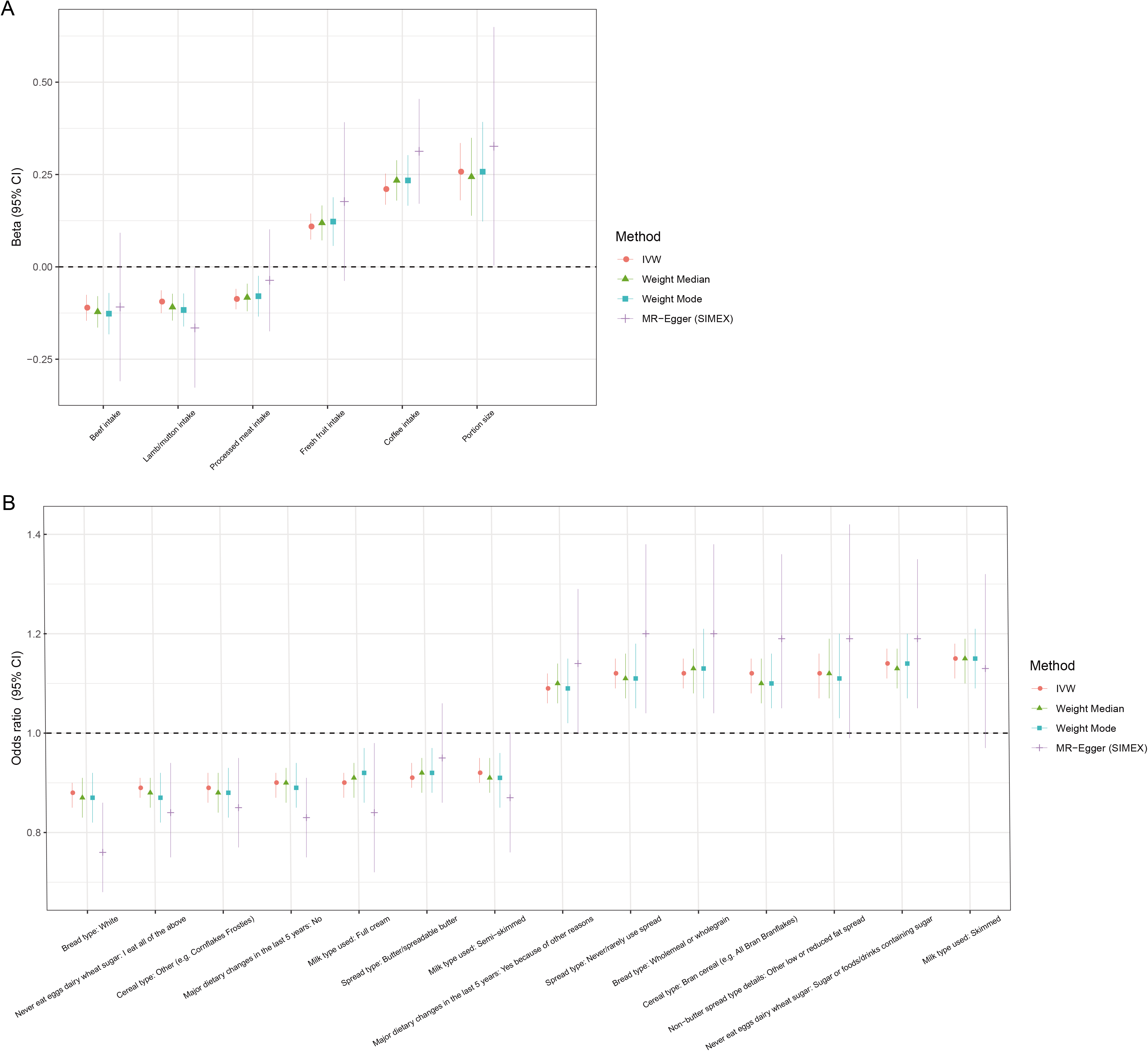
Summary Mendelian randomization (MR) estimates derived from the main inverse-variance weighted (IVW), MR-Egger, weighted median and weighted mode-based methods for dietary habits. A: quantitative trait; B. qualitative trait.

**S9 Fig.**
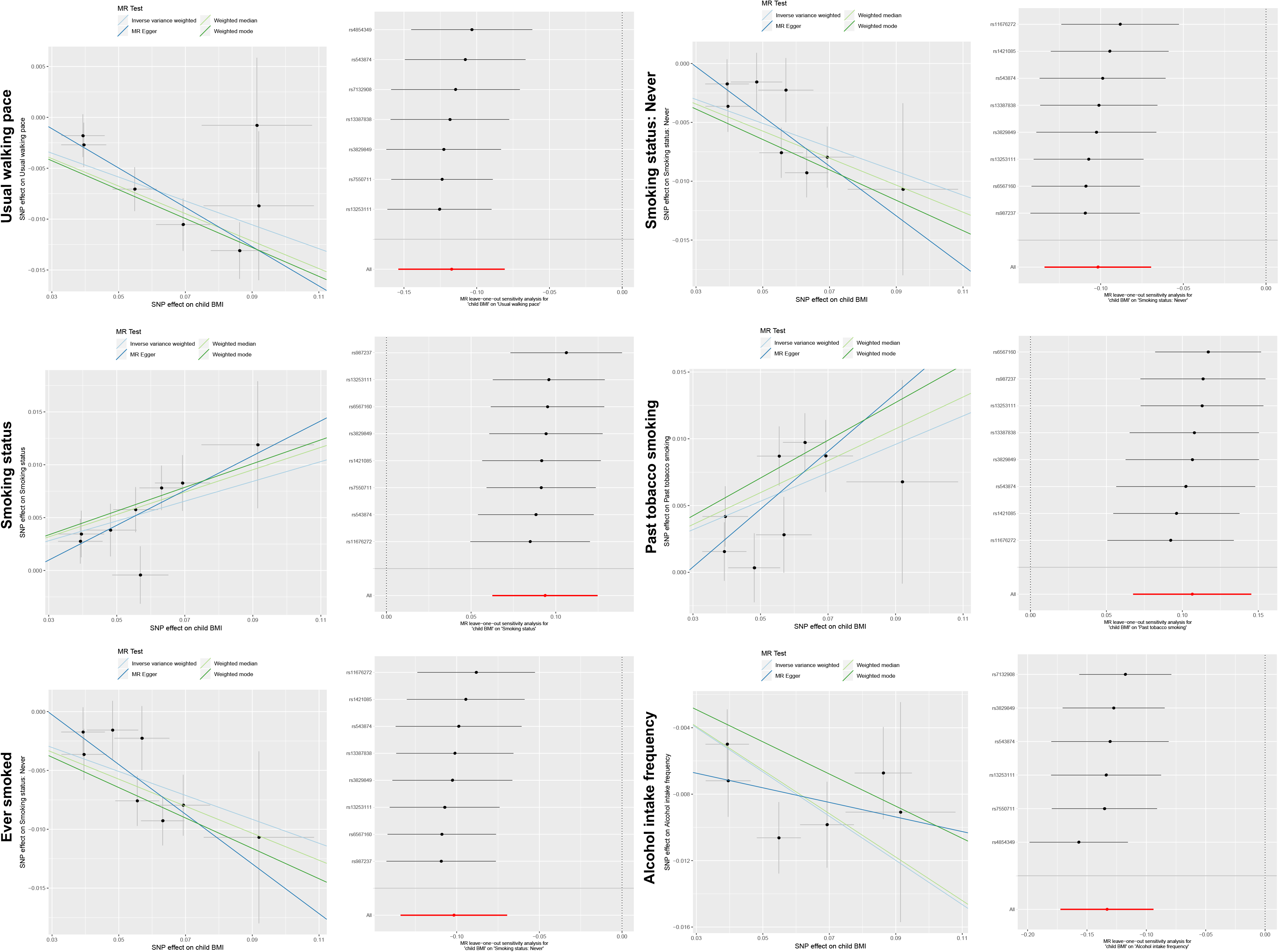
Scatter plot and leave one out analysis plot for physical activity, smoking behaviors, and alcohol intake frequency.

S1 Table. The 53 GWAS summary data from other public resources rather than the UK Biobank population.

S2 Table. Characteristics of the 15 SNPs associated with childhood BMI.

S3 Table. Other studies supporting the associations between 11 of the 13 loci for MR analysis and childhood obesity.

S4 Table. Characteristics of the 76 SNPs associated with adult BMI.

S5 Table. Genetic correlation results. Only 269 outcomes with *P* < 0.05 are shown.

S6 Table. Pleiotropy assessment results for the 269 outcomes. Outcomes using MR-Egger as the main method are marked in yellow.

S7 Table. MR analysis results for the 269 outcomes. 60 outcomes with significant associations with childhood BMI are shown in bold.

S8 Table. MR-analysis using the 4 SNPs not in LD with adult BMI variants for the 60 outcomes associated with childhood BMI. Outcomes with main MR method *P* < 0.05 are shown in bold.

S9 Table. MR-analysis using adulthood BMI as exposure for the 60 outcomes associated with childhood BMI. 76 SNPs were used as instruments.

S10 Table. MR-analysis using adulthood BMI as exposure for the 60 outcomes associated with childhood BMI. 65 SNPs were used as instruments (11 SNPs exists in or in LD with the childhood BMI SNPs were excluded).

S11 Table. Genetic correlation analysis results between the 27 lifestyle factors/socioeconomic status and CAD/T2D. Only results with *P* < 0.05 are shown.

S12 Table. MR analysis results using lifestyle factors/socioeconomic status as exposures and T2D as outcome. Only results with main MR method *P* < 9.26 × 10^−4^ are shown.

S13 Table. Reverse MR analysis results using T2D as exposure and the 7 lifestyle factors as outcome.

